# Inhibition of proline tyrosine kinase 2 (Pyk2) phosphorylation during adherent-invasive *Escherichia coli* infection inhibits intra-macrophage replication and inflammatory cytokine release

**DOI:** 10.1101/2022.11.14.516411

**Authors:** Xiang Li, Michael J. Ormsby, Ghaith Fallata, Lynsey M. Meikle, Damo Xu, Daniel M. Wall

## Abstract

Adherent-invasive *Escherichia coli* (AIEC) have been implicated in the aetiology of Crohn’s Disease (CD). They are characterized by an ability to adhere to and invade intestinal epithelial cells, and to replicate intracellularly in macrophages resulting in inflammation. Proline-rich tyrosine kinase 2 (PYK2) has previously been identified as a risk locus for inflammatory bowel disease through genome wide association studies. It is overexpressed in patients with colorectal cancer, a major long-term complication of CD. Here we show that Pyk2 levels are significantly increased during AIEC infection of murine macrophages while an inhibitor of Pyk2, PF-431396 hydrate, significantly decreased intramacrophage AIEC numbers. Imaging flow cytometry indicated that Pyk2 inhibition blocked intramacrophage replication of AIEC with no change in the overall number of infected cells, but a significant reduction in bacterial burden per cell. This reduction in intracellular bacteria resulted in a 20-fold decrease in tumour necrosis factor α secretion by cells post-AIEC infection. These data demonstrate a key role for Pyk2 in modulating AIEC intracellular replication and associated inflammation and may provide a new avenue for future therapeutic intervention in CD.

## Introduction

Crohn’s disease (CD) is characterized by chronic inflammation of the gastrointestinal tract, most commonly affecting the terminal ileum and proximal colon. A histopathological hallmark of CD is the aggregation of intestinal macrophages, resulting in lesions, referred to as granulomas (1, 2). The initial trigger of CD remains unknown, but the aetiology of CD involves environmental factors, infectious agents, and genetic predisposition. Together these lead to an abnormal mucosal immune response to pathogenic insult and alterations in both microbial composition and metabolic activity of intestinal communities (3). Dysbiosis in CD patients is observed as an overall reduction in gut microbial diversity and an overgrowth of pro-inflammatory bacteria including *Enterobacteriaceae*. In particular, a pathovar of *Escherichia coli* known as adherent-invasive *E. coli* (AIEC) increases in prevalence during inflammation and has been detected in the ileal mucosa of CD patients at higher levels compared to healthy controls (21.7% - 36.4 % versus 6.2%) (4–6). Other studies have directly implicated AIEC in the aetiology of CD (6, 7).

Once AIEC cross the intestinal epithelium, they are phagocytised by macrophages into a double-membraned degradative organelle, termed a phagolysosome (8), that is equipped with a broad range of lytic enzymes (9, 10). AIEC have acquired several factors that are essential for evading phagolysosome degradation; including tolerance of acidic pH and nutritional and oxidative stress (11). However, the ability of AIEC to survive within the harsh environment of a macrophage with the concomitant release of tumour necrosis factor α (TNFα) is still poorly understood.

Intestinal macrophages enriched in the lamina propria, capture and eliminate any bacteria that cross the epithelial barrier and are responsible for clearing apoptotic and senescent epithelial cells (12). These sub-epithelial macrophages are essential for maintenance of mucosal homeostasis in the presence of the microbiota and also play a pivotal role in protective immunity against pathogens (13). Macrophages mount their response against microbial pathogens through binding of pattern recognition receptors (PRRs) to pathogen-associated molecular patterns (PAMPs) resulting in release of a variety of proinflammatory cytokines and chemoattractants, such as TNF α. This cytokine is a key mediator of inflammation in CD, disrupting epithelial barrier function by altering the structure and function of tight junctions (14, 15). A milestone in treating CD was the introduction of anti-TNFα agents, like infliximab and adalimumab (16, 17).

Large-scale genome-wide association studies in cohorts of European patients resulted in the identification of candidate genes in inflammatory bowel disease (IBD) susceptibility loci (18). These included proline-rich tyrosine kinase 2 (*PYK2*), a non-receptor, Ca^2+^ dependent protein-tyrosine kinase that is expressed in numerous tissues and cell types and which is involved in innate immunity (19). It is highly expressed in the central nervous system, epithelial cells, hematopoietic cells, and it is also over-expressed in various cancers (20, 21). Activation of PYK2 involves autophosphorylation at Tyr-402, which enables the binding of Src via the SH2 domain and phosphorylation of PYK2 at Tyr-579 and Tyr-580, within the kinase domain activation loop, to generate maximal kinase activity (22). Phosphorylated PYK2 functions in the regulation of phagocytosis, migration, proliferation, invasion, oncogenesis and metastasis (21, 23–25). In macrophages, PYK2 has defined functions in regulating morphology, migration, and phagocytosis (21, 26, 27). It is therefore considered a valuable therapeutic target in various disease states, such as inflammation and cancer (28).

In this study, the role of Pyk2 in facilitating intracellular replication of the AIEC type-strain LF82 in murine macrophage cells was determined. Through the use of high throughput imaging of individual cells via imaging flow cytometry, we have gained an increased understanding of the role of this kinase at the single cell level, demonstrating a key role for Pyk2 in controlling intramacrophage replication of this poorly understood pathogen.

## Material and methods

### Bacterial strains and culture conditions

The adherent-invasive *E.coli* (AIEC) type strain LF82 used in this study was cultured in lysogeny-broth (LB) at 37°C with shaking at 180 revolutions per minute (rpm) (29). Prior to infection, LF82 was grown overnight in Roswell Park Memorial Institute (RPMI)-1640 media (Life Technologies) supplemented with 3% heat-inactivated foetal bovine serum (FBS; Sigma), and 1% L-glutamine. *In vitro* infections were carried out using LF82 transformed with a reporter plasmid, pAJR70 expressing eGFP under the control of the *rpsM* promoter (30). GFP expression was measured in a black 96-well plate (excitation at 485 nm; emission at 550 nm) using a FLUOstar Optima Fluorescence Plate Reader (BMG Labtech).

### Maintenance of RAW 264.7 cells

The RAW 264.7 murine macrophage-like cell line was obtained from the American Type Culture Collection (ATCC). RAW 264.7 cells were cultured in RPMI-1640 media supplemented with 10% heat-inactivated FBS, 1% penicillin/streptomycin, and 1% L-glutamine (maintenance media) at 37°C in 5% CO_2_.

### Gentamicin protection assay for AIEC infection

RAW 264.7 cells were seeded onto a 24-well plate at a density of 2×10^5^ cells-per-well 24 h prior to infection (hpi). Twelve hours prior to infection, maintenance media was replaced with RPMI-1640 media supplemented with 3% FCS and 1% L-glutamine containing lipopolysaccharide (LPS 1 μg/ml) for activation. Cells were infected at a multiplicity of infection (MOI) of 100 and incubated at 37°C/5% CO_2_ for 1 h. After 1 h the infected RAW 264.7 cells were washed twice with fresh RPMI cell culture media to remove excess extracellular bacteria and incubated at 37°C/5% CO_2_ in maintenance media with 50 μg/ml gentamicin to kill any remaining extracellular bacteria. In addition to gentamicin, cell culture media was also supplemented where indicated with the Pyk2 inhibitor PF-431396-hydrate (31). The infected cells were then incubated for the time specified at 37°C/5% CO_2_.

To measure AIEC intra-macrophage survival, infected macrophages were washed with RPMI-1640 media and lysed using 200 μl of 2% Triton X-100 in phosphate buffered saline (PBS) for 5 min at room temperature. Lysates were removed, serially diluted in PBS, and plated onto LB agar plates to determine the number of colony forming units (CFU) per mL. Total protein concentration was determined using a BCA assay (Thermofisher) and bacterial numbers were normalised to total protein concentration and presented as CFU/g. Normalizing CFU to protein concentration, as opposed to expressing CFU numbers per well, meant CFUs could be related to cell number in each individual well, especially important if cell numbers differed between wells due to proliferation or cell death during infection or drug treatment.

### AIEC phagocytosis assay

RAW 264.7 cells were plated in a 24-well plate at a density of 2×10^5^ cells-per-well. Cells were incubated (37°C, 5% CO2) for 6 h resulting in adherence to plates. Before infection, cells were activated by adding 1 μg/ml of LPS 12 h prior to infection. Activated cells were treated with the indicated concentrations of PF-431396 hydrate (Sigma-Aldrich) for 1 h. PF-431396 hydrate suppresses the phosphorylation of Pyk2 at its active tyrosine phosphorylation site, Y402, without decreasing total protein level of Pyk2, thus functionally blocking multiple cellular signalling pathways (32). Supernatants were removed and cells were infected with an MOI of 100 in 200 μl of RPMI-1640 media containing 3% FCS and 1% L-glutamine. Cells were left for 1 h to allow bacterial phagocytosis to occur, before the supernatant was removed and 1 ml of fresh media containing gentamicin (50 μg/mL) was added for 1 h to kill any extracellular bacteria. Post-gentamicin treatment, supernatants were removed, cells were lysed with 200 μl PBS containing 2% Triton X-100 for 5 min and bacteria numbers calculated as described above.

### Lactate dehydrogenase assay to monitor cytotoxicity

RAW 264.7 cells were infected at an MOI of 100 and supernatants of LF82 infected and control macrophages were sampled at different time points of gentamicin treatment, centrifuged at 10,000 x g for 4 min and assayed for lactate dehydrogenase (LDH) activity (Sigma). LDH activity is reported as milliunit/ml. One unit of LDH activity is defined as the amount of enzyme that catalyses the conversion of lactate into pyruvate to generate 1 μmol of NADH per minute at 37°C.

### Cell counting kit 8 (CCK-8) assay for detection of cell viability

Cell survival rates were estimated by the CCK-8 assay (Abcam). Approximately 10^4^ cells were seeded in 96-well plates with 100 μl medium each well. After 24 h cultivation, different doses of PF-431396 were added for a further 6 h. Each well was incubated with 10 μl of CCK-8 solution for 2 h away from light before measuring the absorbance at 450 nm by FluoStar Optima fluorescent plate reader (BMG Biotech). The relative viability was expressed by the formula: % of viability = ((A_exp_ – A_Blank_)/A_control_ - A_Blank_)) X 100%.

### Western Blotting

LF82 infected and control cells were lysed at 6 hours post-infection (hpi) for 15 min in radioimmunoprecipitation assay (RIPA) lysis buffer (ThermoFisher) supplemented with protease inhibitor (Sigma) and phosphatase inhibitor (Sigma). Lysates were centrifuged at 14,000 x g for 10 min. The protein concentration of the cell extracts was determined using the BCA assay kit (Bio-Rad). Protein concentration for lysates was adjusted to 2 μg/μl before adding 4X loading buffer (ThermoFisher), heating to 95 °C for 10 min and running on an SDS-PAGE gel. Samples were transferred via electrophoretic wet transfer to a polyvinylidene fluoride (PVDF) membrane. The membrane was blocked in 5% non-fat milk and probed with a 1:1,000 dilution of anti-Pyk2 (Abcam) or a 1:1000 dilution of anti-phosphorylated Pyk2 (ab4800; Abcam). Blots were visualised with HRP-conjugated goat anti-rabbit antibody (1:10,000) or HRP-conjugated goat anti-mouse (1:10,000) (ThermoFisher) and developed using enhanced chemiluminescence (ECL) and imaged using a C-DiGit blot scanner (LI-COR). Membranes were stripped in 0.1 mM glycine, pH 2.2, and re-probed with anti-GAPDH (1:10,000; Abcam) antibody as a loading control. The bands were quantified using ImageJ software (33). To compare both Pyk2 and pPyk2 expression in infected or uninfected RAW 264.7 cells with or without PF-431396 hydrate treatment, values from each blot were normalised by loading control (GAPDH) as Pyk2/GAPDH or pPyk2/GAPDH. The Pyk2/GAPDH and pPyk2/GAPDH values of control groups were set as 1 and the relatively densitometric change of test samples was calculated. All Western blots were performed in triplicate, with each performed on a biological replicate. All data are represented as the mean ± standard deviation for all performed repetitions.

### Immunofluorescence

Images for all experiments were captured using a Leica DMi8 fluorescent microscope. RAW 264.7 Cells were plated onto glass coverslips in 24-well-plates at 2.0×10^5^ cells per well. Cells were infected with LF82::*rpsM*GFP for 1 h then either treated with the Pyk2 inhibitor PF-431396 hydrate, or left untreated, for 6 h or 12 h. At each time point, cells were fixed immediately with 4% paraformaldehyde (PFA) solution and permeabilized with 0.2% Triton X-100 after rinsing with Dulbecco’s Phosphate-Buffered Saline (DPBS). Coverslips were treated with 1% bovine serum albumin (BSA)/DPBS to block non-specific binding for 30 min at 37°C. The following primary and secondary antibodies were used: anti-pPyk2 (ab4800; Abcam) followed by Alexa Fluor® 647 conjugated goat anti-rabbit antibody (ThermoFisher). Phalloidin (ThermoFisher) and DAPI (VECTOR) were used for visualizing F-actin and the nucleus, respectively.

### Imaging flow cytometry

Imaging flow cytometry (IFC) data acquisition was achieved using an ImageStream X MKII (Amnis) equipped with dual cameras and 405 nm, 488 nm, and 642nm excitation lasers. All samples were acquired at 60 times magnification giving an optimal 7 μM visual slide through the cell, and a minimum of 10,000 single cell events were collected for each sample. In focus cells were determined by a gradient root mean square (RMS) for image sharpness. Brightfield of greater than 50 and single cells were identified by area versus aspect ratio. Only data from relevant channels were collected including Channel 02 (Ch02, GFP fluorescence), Ch04 (bright field), and Ch05 (pPyk2 conjugated AF647 fluorescence). Samples were run with a 488 nm laser with power of 5 mW and a 642 nm laser with power of 150 mW. Single colour compensation controls were also acquired, and a compensation file generated via IDEAS software (Luminex).

### Imaging flow cytometry data analysis

Data analysis for samples collected using the ImageStream X MKII was performed using IDEAS. Masks (area of interest) and features (calculations made from masks) were generated to give quantitative measurements of the images collected. To quantify intracellular bacteria, a mask was created to select just the intracellular portion of the cell, in this case referred to as an AdaptiveErode mask. To analyse fluorescent spots (representative of bacterial burden per cell), a spot count mask was created in the reference channel for GFP bacteria (Ch02).

### Enzyme-linked immunosorbent assays (ELISA)

The amount of TNFα secreted in the supernatants from cell culture and cell lysate was determined by ELISA MAX™ Deluxe Set Mouse TNFα kit (BioLegend) according to the manufacturer’s instructions. TNFα concentrations were normalised by protein concentration from cell lysates and were reported as ρg of TNFα/ g of protein.

### Statistical analysis

Values are shown as means and standard deviation. All statistical tests were performed with GraphPad Prism software, version 8.3.0. All replicates in this study were biological; that is, repeat experiments were performed with freshly grown bacterial cultures and cells, as appropriate. Technical replicates of individual biological replicates were also conducted. Significance was determined as indicated in the figure legends. Values were considered statistically significant when *p*-values were ∗ = *p*<0.05; ∗∗ = *p*<0.01; ∗∗∗ = *p*<0.001; ∗∗∗∗ = *p*<0.0001.

## Results

### Macrophage Pyk2 levels increase in response to L82 infection

To determine the change in Pyk2 expression in RAW 264.7 macrophages in response to LF82 infection, total Pyk2 and phosphorylated Pyk2 (pPyk2) levels were measured by immunoblotting (Fig. 1). A significant increase in total Pyk2 was observed at 6 hours post-infection (hpi) and pPyk2 levels were also increased (Fig. 1a-b). To investigate this further in relation to infection, imaging flow cytometry was used to analyse pPyk2 levels simultaneously in thousands of individual uninfected and LF82-infected cells. Imaging flow cytometry allows identification of all cells within a population that are infected with GFP-expressing bacteria. These cells can be counted and the individual bacterial load in each determined, allowing the separation of cells based on both their infection status and bacterial load in a way not possible with traditional colony-counts. Using this approach here it was determined that there was a significant increase in pPyk2 levels in response to LF82 infection (Fig. 1c-d).

**Figure 1:**
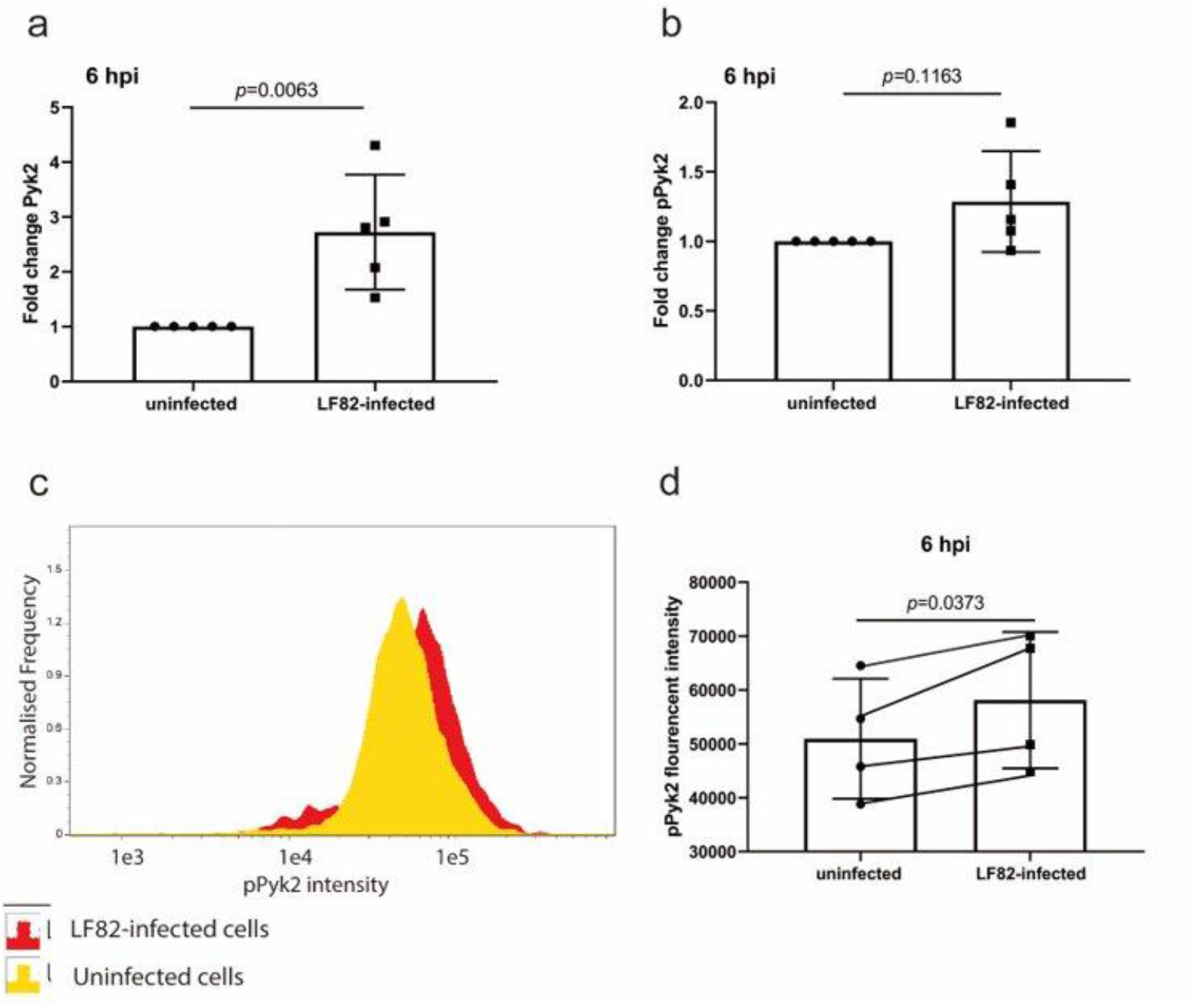
Pyk2 and phosphorylated Pyk2 (pPyk2 [Y402]) expression levels in RAW 264.7 macrophages. (**a**) A representative Western blot at 6 hpi is shown. Levels of Pyk2 (**b**) or pPyk2 (Y402) (**c**) at 6 hpi were compared between LF82-infected or uninfected RAW 264.7 cells using densitometric analysis of Western blots. Bands from infected cells (**a**) were analysed by densitometry and compared to control uninfected cells using ImageJ and fold change in Pyk2 (**b**) and pPyk2 (Y402) levels (**c**) are shown. Statistical analyses were conducted using a Student’s t-test. (**d**) A representative histogram showing imaging flow cytometry data displays the pPyk2 (Y402) intensity of control uninfected RAW 264.7 macrophages in yellow and LF82 infected cells at 6hpi in red. (**e**) The level of pPyk2 (Y402) in infected RAW 264.7 cells at 6 hpi was analysed by imaging flow cytometry and compared with that of uninfected cells, using a paired t-test statistical analysis. Results are representative of at least four independent experiments.

### Pyk2 inhibitor PF-431396 hydrate successfully blocks phosphorylation of Pyk2 in uninfected and LF82 infected macrophages

PF-431396 hydrate inhibits phosphorylation of Pyk2 at its active site Y402, blocking multiple cellular signalling pathways (32). Activated Pyk2 was measured by immunoblotting cell extracts with phospho-specific antibodies directed against the Pyk2 (Y402) autophosphorylation site. Six hours post-infection levels of Pyk2 or pPyk2 (Y402) were measured by Western blot in infected cells and controls, including those where PF-431396 hydrate had been added (Fig. 2) PF-431396 hydrate significantly reduced levels of pPyk2 (Y402) in LF82-infected and uninfected RAW 264.7 cells post-treatment (5 and 10 μM; Fig.2a).

**Figure 2:**
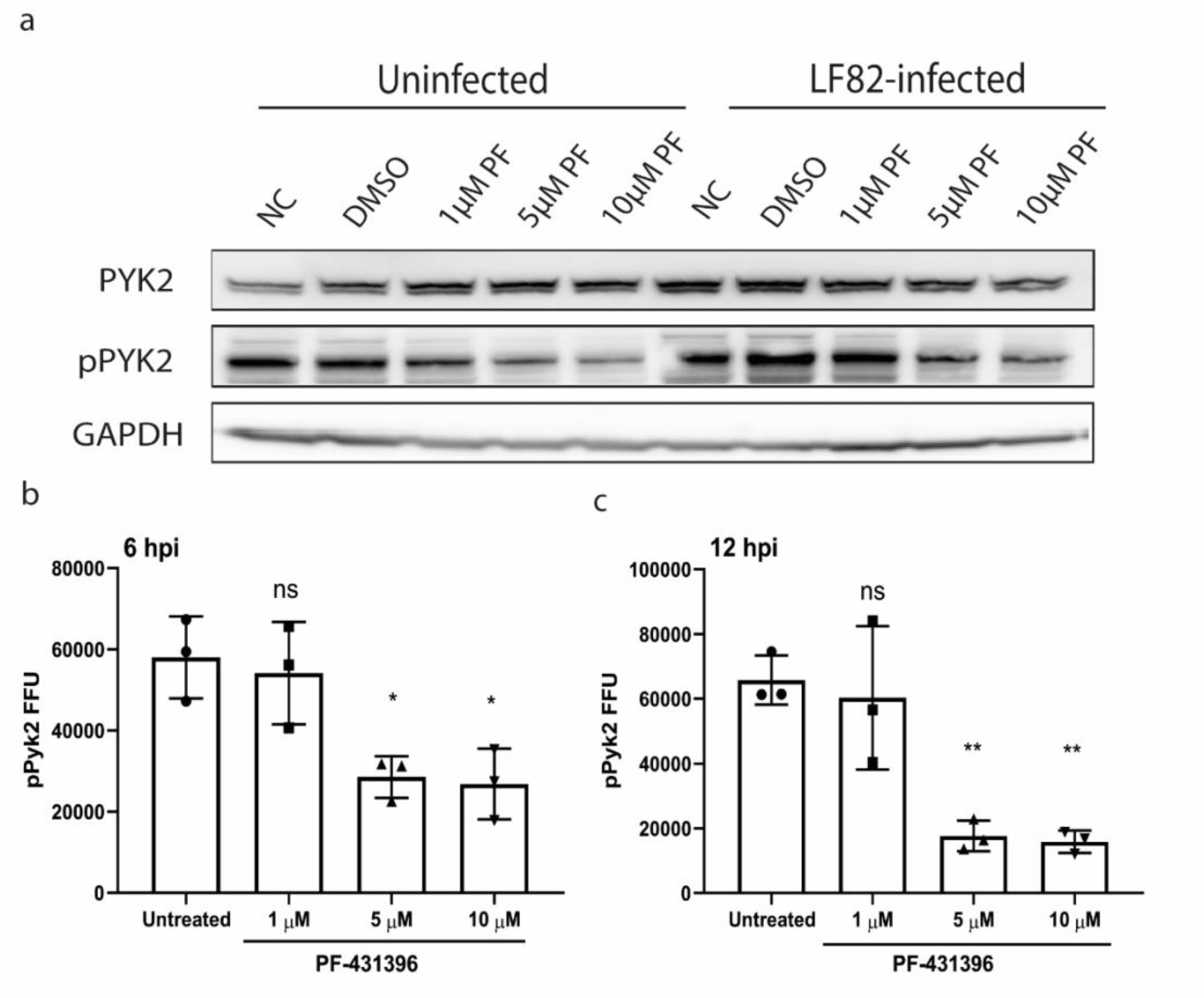
The inhibitor PF-431396 hydrate blocks phosphorylation of Pyk2 in both uninfected and LF82 infected cells in a dose dependent manner. LF82 infected or control uninfected RAW 264.7 cells were treated with PF-414396 hydrate (0, 1, 5, 10 μM) for 6 h. Untreated cells were used as controls. (**a**) Immunoblotting was used to detect levels of pPyk2 (Y402) with GAPDH used as loading control. To determine relative pPyk2 (Y402) levels in inhibitor treated cells, imaging flow cytometric analysis was conducted on LF82 infected RAW 264.7 cells after treatment for 6 h (**b**) or 12 h (**c**) with PF-431396 hydrate. Statistical analyses were conducted using a one-way ANOVA test. (ns, not significant, \**p* < 0.05, \*\**p* < 0.01, \*\*\**p* < 0.001). Data are representative of three independent biological replicates.

Imaging flow cytometric analysis further confirmed that PF-431396 hydrate significantly reduced pPyk2 (Y402) levels in infected cells with the number of cells with detectable pPyk2 (Y402) significantly dropping at 6 hpi (Fig. 2b) before reducing further at 12 hpi (Fig. 2c). To ensure that these decreases in pPyk2 (Y402) levels were not due to cytotoxicity of PF-431396 hydrate, we demonstrated no increased release of lactate dehydrogenase (LDH) in uninfected or infected RAW 264.7 cells when PF-431396 hydrate was used at concentrations up to, and including, 10 μM (Fig. S1).

### Pyk2 is important for intracellular survival of AIEC in macrophages

Given the previously reported role of Pyk2 in phagocytosis (27), we analysed the role of Pyk2 in phagocytosis of LF82 by RAW 264.7 cells. RAW 264.7 cells were infected with LF82 following a 1 h pre-treatment with PF-431396 hydrate before bacterial colony forming units (CFUs) were counted. The highest concentration of inhibitor (10 μM), significantly impaired phagocytosis compared to cells that were untreated or treated with a lower concentration of inhibitor (Fig. 3a). Having established that Pyk2 plays a role in phagocytosis of LF82, further treatments to inhibit Pyk2 were conducted post-phagocytosis of LF82. This was to ensure that any alteration in intracellular numbers of LF82 were not impacted by the inhibition of phagocytosis.

**Figure 3:**
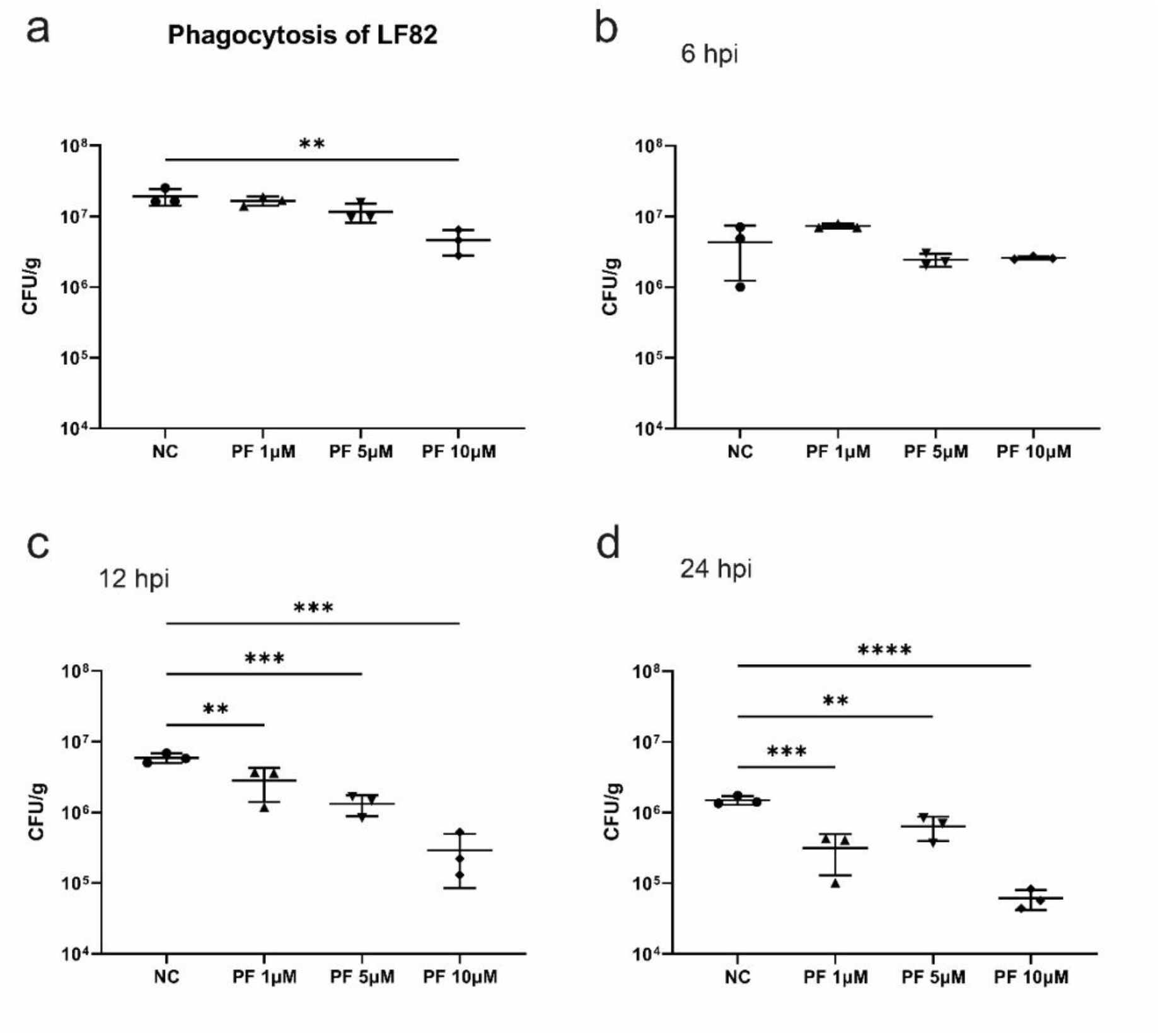
Pyk2 inhibition reduces intracellular LF82 in RAW 264.7 cells. (**a**) (**a**) To determine its effect on phagocytosis of LF82, RAW 264.7 cells were pre-treated with PF-431396 hydrate for 2 hours before infection with LF82 at an MOI of 100. PF-431396 hydrate was added to RAW 264.7 cells at concentrations of 0 μM (NC – negative control), 1 μM, 5 μM or 10 μM. DMSO in which PF-43196 hydrate was initially aliquoted was added as an additional control to ensure no residual effects of its presence on Pyk2 phosphorylation. Intracellular replication of LF82 in the presence of PF-431396 was measured by total viable counts at (**b**) 6 hpi, (**c**) 12 hpi and (**d**) 24 hpi. PF-431396 hydrate was added to RAW 264.7 cells at concentrations of 0 μM (NC – negative control), 1 μM, 5 μM or 10 μM 1 hpi. Statistical analyses were conducted using a one-way ANOVA. (ns, not significant, \**p* < 0.05, \*\**p* < 0.01, \*\*\**p* < 0.001, \*\*\*\**p* < 0.0001). Data are representative of three independent biological experiments.

### Inhibition of Pyk2 function significantly reduces intra-macrophage LF82 burden

Intracellular replication is a hallmark of the AIEC phenotype. To determine the role of Pyk2 in intracellular replication of LF82, PF-431396 hydrate was added at concentrations of 0, 1, 5 or 10 μM at 1 hpi. Intracellular bacteria numbers were measured using colony counts at 6, 12 and 24 hpi. At 6 hpi, there was no significant difference in intracellular bacterial numbers relative to the control at any concentration of PF-431396 hydrate (Fig. 3b). However, a significant reduction in intracellular LF82 numbers was seen at 12 hpi with both 5 μM and 10 μM PF-431396 hydrate (Fig. 3c), and at 24 hpi for all concentrations of PF-431396 hydrate (Fig. 3d).

The interaction of intracellular pathogens with host cells is traditionally quantitated on a cellular population level. However, intracellular replication is heterogeneous, and infection of host cells with a clonal population of any intracellular pathogen frequently results in variable numbers of bacteria in individual host cells. To examine this, immunofluorescence was conducted to permit direct visualisation of infected cells. In addition, imaging flow cytometry was conducted to allow quantification of morphological cellular features and spatial distribution of fluorescent markers at the single cell level in the heterogenous infected cell population. This made it possible to correlate acquired cellular images with events and accurately quantify intracellular bacteria.

To establish that transformation of LF82 with the fluorescent *rpsM*GFP reporter required for intracellular imaging did not alter the bacterial phenotype, growth rates of LF82 and LF82::*rpsM*GFP were assessed in the presence and absence of PF-431396 hydrate. There were no significant differences between any of the conditions assessed (Fig. S2a&b). An intracellular infection model, where the output is viable bacterial counts, was used to assess bacterial replication and survival in RAW 264.7 macrophages transformed with LF82::*rpsM*GFP at 6, 12 and 24 h (Fig. S2c-e). Comparison of intracellular replication of LF82 and LF82::*rpsM*GFP at 6, 12 and 24 hpi demonstrated that there was no significant difference in intracellular survival or replication between the strains (Fig. S2c-e), making LF82::*rpsM*GFP a suitable fluorescent reporter strain.

### Pyk2 inhibition blocks LF82 replication intracellularly without affecting overall numbers of infected cells

Given that LF82 infection of RAW 264.7 cells is heterogenous with potentially large numbers of intracellular bacteria, imaging flow cytometry was employed to quantitatively correlate the number of intracellular bacteria with the level of Pyk2 phosphorylation and activation in single cells. Single cells were gated as described in Fig. S3. Utilising IDEAS imaging flow analysis software (Luminex) spot count masks were created and used in conjunction with the erode mask to produce intracellular spot count profiles for each time point (Fig S4a-d).

To facilitate analysis cells were assigned to groups with those with greater than 10 bacteria deemed to have a “high” bacterial burden, those with 6-10 bacteria assigned to an “intermediate” bacterial burden, and those with five or less as having a “low” bacterial burden. While assigning cells to these groups was arbitrary, cells were assigned to groups based on the hypothesis that cells with greater than 10 bacteria were more likely to be representative of those where intracellular replication of LF82 has taken place, whilst those with fewer than five bacteria were more likely to be indicative of bacterial phagocytosis rather than replication. Separating the infected macrophage population into distinct populations based on bacterial load is a significant step forward in understanding the dynamics of LF82 infection, which was previously determined using highly heterogenous populations of infected cells. For each of high, intermediate, and low bacterial burdens the percentage of the total population was plotted at 6 hpi (Fig. 4a) and 12 hpi (Fig. 4b). Since treatment with the Pyk2 inhibitor was initiated post-phagocytosis, and therefore did not influence bacterial uptake by the macrophages, there were no significant differences observed in the ratio of infected:uninfected cells within any of the treatment groups. Untreated cells or those treated with 1 μM of PF-431396 hydrate had a significantly larger population of cells with high bacterial burden at both 6 hpi (18.57% and 16.96% of total cells respectively) and 12 hpi (17.56% and 17.26% respectively) compared to 10 μM treatment, which resulted in a significant decrease in the number of cells with a high bacterial burden (9.01% at 6 hpi and 6.87% at 12 hpi, Fig.4a and 4b). In cells treated with 10 μM PF-431396 hydrate, where less pPyk2 (Y402) protein was detectable, a significantly larger population of cells were observed to have a low bacterial burden at both 6 hpi (49.13%) and 12 hpi (53.43%), compared to untreated cells which had 39.65 % and 41.53 % of cells deemed to have a low bacterial burden. This expansion of low bacterial burden cells in 10 μM PF-431396 hydrate treated cells was at the expense of those with a high bacterial burden, as there was a corresponding reduction in this subset. These data indicate that when phosphorylation of Pyk2 is inhibited it has a direct inhibitory effect on intracellular replication of LF82 in RAW 264.7 macrophages.

**Figure 4:**
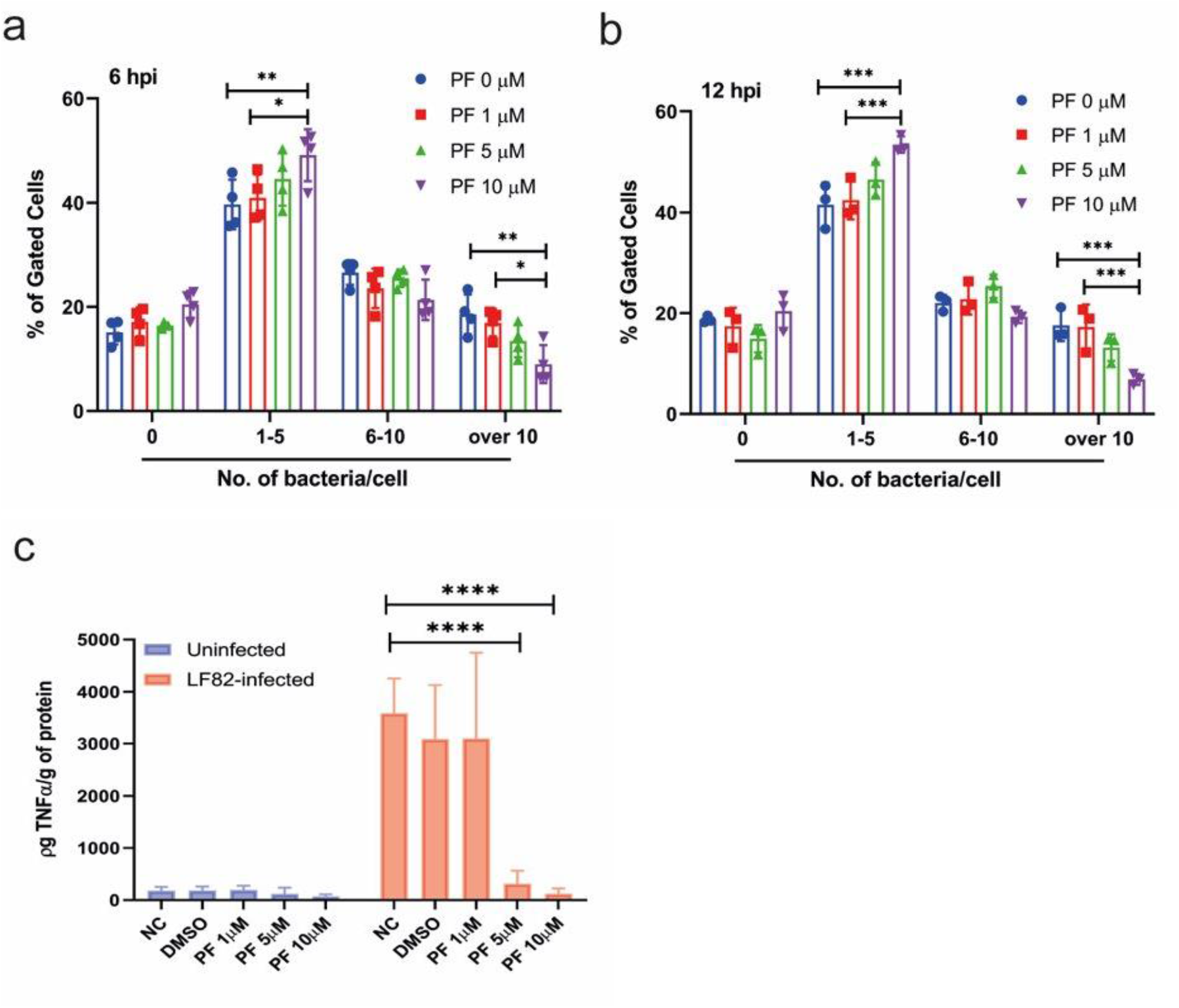
Treatment with PF-431396 hydrate inhibits Pyk2 phosphorylation and reduces intra-macrophage LF82 burden and TNFα secretion. LF82::*rpsM*GFP burden in RAW 264.7 macrophages was analysed via imaging flow cytometry. The bacterial spot count profiles at 6 hpi (**a**) and 12 (**b**) hpi were separated into: groups representing uninfected cells (0 bacterium); cells with a low infection count (1-5 bacteria per cell); medium infection count (6-10 bacteria per cell); and highly infected cells (>10 bacteria per cell) where PF refers to concentration of PF-431396 hydrate added. Statistical analysis was conducted using a two-way ANOVA test. (ns, not significant, \**p* < 0.05, \*\**p* < 0.01, \*\*\**p* < 0.001, \*\*\*\**p* < 0.0001). Data are representative of three independent biological replicates. (c) Comparison of TNF-α section levels from supernatants of infected or uninfected RAW 264.7 cells. RAW 264.7 cells were activated by LPS overnight then infected or uninfected with LF82 for 1 h followed by treatment with PF-431396 hydrate for 6 h. At 6 hpi, TNF-α levels in supernatants were examined by ELISA. Values are mean ± SD (n = 3 per group). Statistical analysis was conducted using a two-way ANOVA test (ns, not significant, *p < 0.05, **p < 0.01, ***p < 0.001, ****p < 0.0001).

### PF-431396 hydrate-mediated blocking of LF82 replication in macrophages significantly reduces TNFα secretion

High levels of TNFα are detected in CD patients and secretion of TNFα occurs post-infection of macrophages by AIEC (34). High levels of TNFα are released as consequence of mucosal injury and transmural inflammation in CD, primarily from lamina propria mononuclear cells (35). Additionally previous studies have highlighted a role for Pyk2 activation in TNFα secretion (36, 37). Given intracellular numbers of LF82 were directly affected by inhibition of PyK2 using PF-431396 hydrate, we went on to determine if there was any downstream impact on TNFα release. Treatment of macrophages with 5 μM and 10 μM PF-431396 induced a significant reduction in TNFα release (between 15- and 40-fold) (Fig.4c). This reduction was not seen in untreated macrophages or those treated with 1 μM inhibitor (Fig.4c). These decreases in TNFα release could not be attributed to increased toxicity due to the inhibitor (Fig. S1). While it is difficult to conclusively conclude that this reduction in TNFα secretion is a direct result of Pyk2 inhibition, rather than an indirect effect of the lower intracellular burden of LF82, the significant decrease in TNFα release that was observed at 5 μM PF-431396 hydrate was not associated with a reduction in intracellular burden (Fig. 4a&b). After establishing that Pyk2 inhibition reduced intra-macrophage replication of LF82, and secretion of TNFα by LF82 infected macrophages, we examined the role of Pyk2 more widely. Similar effects of Pyk2 inhibition were also observed in macrophages infected with *Salmonella enterica* serovar Typhimurium (*S*. Typhimurium) strain SL1344 and clinical isolates of *E. coli* (B94, B115 and B125) from CD patients. PF-431396 hydrate treatment (5 μM) resulted in significantly reduced intracellular bacteria number in macrophages for all clinical isolates except *E. coli* B122 and the commensal *E. coli* F18 strain (Fig. 5a&b). Pyk2 inhibition significantly decreased TNFα secretion in macrophages infected with all AIEC clinical isolates, including strain B122, at both 6, and 24 hpi (Fig. 5c&d). While large reductions in TNFα secretion were seen in macrophages that had taken up SL1344 and *E. coli* F18 at both 6, and 24 hpi, these reductions were not statistically significant.

**Figure 5:**
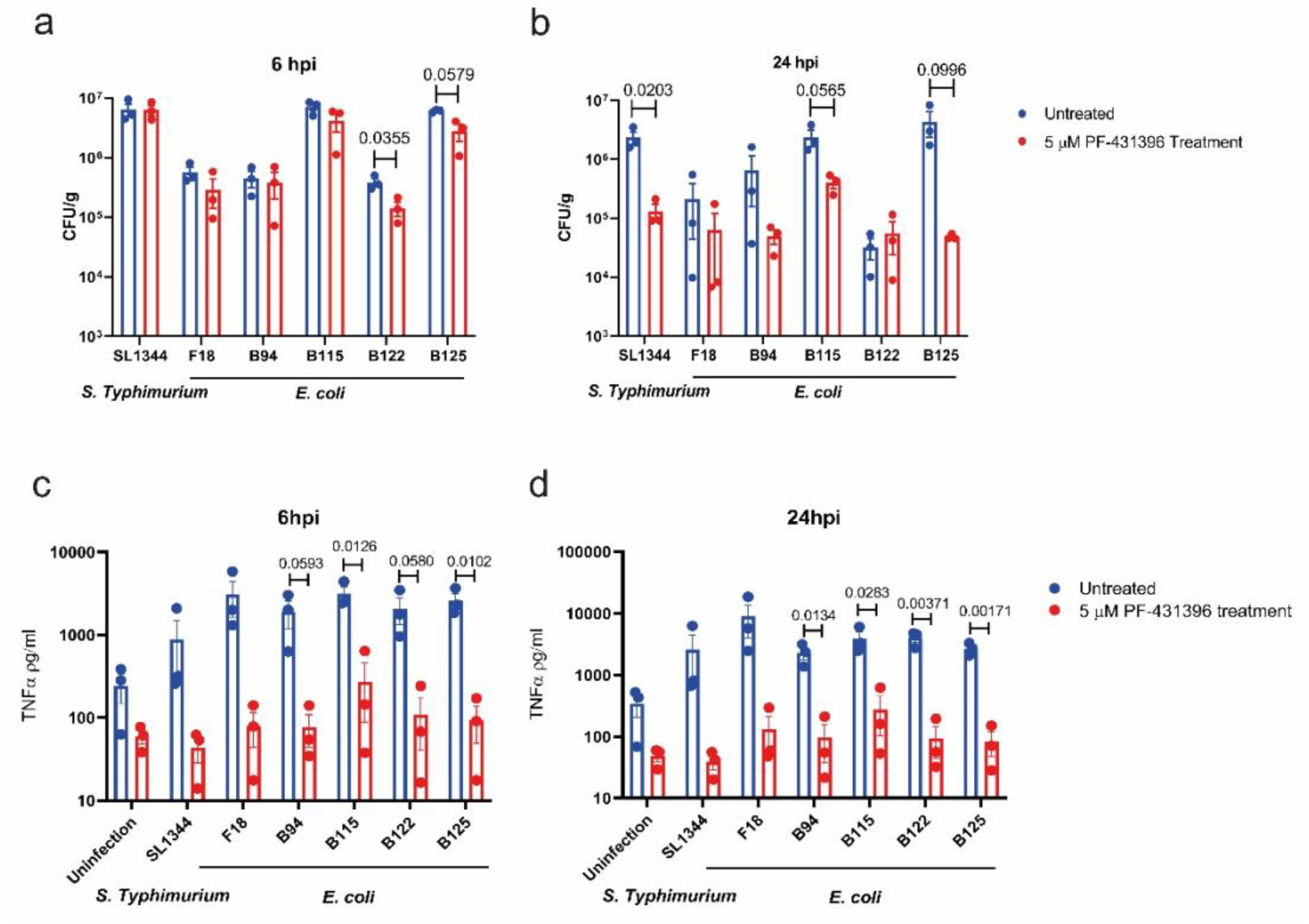
Pyk2 inhibition reduces AIEC clinical isolate intramacrophage burden and TNFα release. RAW 264.7 macrophages were infected with AIEC type strain, LF82, commensal strain *E. coli* F18, *Salmonella* Typhimurium strain SL1344 and AIEC clinical isolates B94, B115, B122, and B125 for 1 hour with MOI of 100, followed by treatment with or without 5 μM PF-431396 hydrate for a further 6, or 24 hpi. (**a-b**) intracellular bacterial numbers were determined by CFU counts. (**c-d**) ELISAs were conducted to detect the secretion of TNFα from uninfected or infected macrophages with or without 5 μM PF-431396 hydrate treatment. Data displayed are of three independent experimental replicates, with each experiment including three independent biological replicates. Data are expressed as mean ± SEM; data were analysed using unpaired multiple t test.

## Discussion

Proline tyrosine kinase 2 (Pyk2) has been identified as a susceptibility locus for IBD risk (18). However, an association between Pyk2 and AIEC, the *E. coli* pathotype consistently overrepresented in the CD intestinal microbiome, has never been investigated. For the first time, our work has demonstrated a direct link between AIEC intracellular replication and activation of Pyk2. Here we show that Pyk2 plays a dual role in AIEC infection, by initially impacting bacterial phagocytosis before subsequently facilitating AIEC replication and survival within macrophages.

To date, only infection by *Y. pseudotuberculosis* has been shown to require Pyk2 for bacterial phagocytosis by macrophages (38, 39). Previous studies identified that the *Yersinia* outer surface protein, invasin, provides high-affinity interactions with host cell surface β1 integrin receptors (40). Binding of integrins then induces integrin clustering and sustained activation of Pyk2, which has been implicated in numerous actin-based cellular processes including cell cycle progression, adhesion and migration (41). Consistent with this, Bruce-Staskal, *et al*. (2002) have shown that *Yersinia* uptake involved a complex interplay in Cas, FAK, Pyk2 and Rac1 cell signalling (42). Through inhibition of Pyk2 phosphorylation, we first demonstrated that AIEC phagocytosis (Fig. 2) was impeded, again reflecting the known role of phosphorylation of Pyk2 in the regulation of phagocytosis (27, 28). We then determined that in addition, the AIEC type strain LF82 is reduced intracellularly in a dose-dependent manner in response to inhibition of Pyk2 function.

Phagocytosis is initiated by macrophage cell membrane Toll-like receptor (TLR) 4 and TLR 5, which recognise bacterial extracellular structures such as fimbriae, flagella, LPS and peptidoglycan (43). Upon phagocytosis of microbial pathogens, TLR4 recruits MyD88 initiating the nuclear factor-κB (NF-κB) signalling cascade (44). Xi *et al.* (2010) showed that Pyk2 interacts with MyD88 and regulates MyD88-mediated NF-κß activation in macrophages with resulting TNFα secretion (45). Secretion of high levels of TNFα are associated with AIEC replication and survival within macrophages without inducing cell death (34). To date, the most effective CD treatments include anti-TNFα or anti-integrin treatments to reduce the circulating concentration of TNFα (46). However, the cellular mechanisms underpinning the effect of TNFα on AIEC replication have remained elusive. *In vivo* infection experiments have demonstrated that CD mucosa-associated *E. coli* killing by macrophages could be inhibited by microbial mannan in a TLR4 and MyD88-dependent manner (47). Taken together, this suggests a mechanism supporting AIEC replication and survival within macrophages via a TLR4-MyD88-Pyk2-NF-κß-TNFα cascade.

Intracellular bacterial replication within macrophages is a characteristic trait of AIEC infection. To investigate this, we analysed intracellular replication using colony counts, immunofluorescence, and imaging flow cytometry. All three methods indicated that inhibition of Pyk2 function significantly decreases intracellular replication of AIEC at 12 hpi compared to untreated cells. While no significant differences in intracellular bacteria were observed using viable counts after 6 h (Fig. 2a), there was a significant change in intracellular bacterial numbers observed using imaging flow cytometry. While this tandem approach gives a greater overall picture of the role of Pyk2 in AIEC infection, the contrasting outputs from each at 6 hpi are noteworthy. Viable counts are most commonly used in infection studies, but they are limited by the need for sample processing and subsequent growth of bacteria before counting. They are also heavily dependent on the successful removal of extracellular bacteria by antibiotics. In such a heterogenous mixture of infected cells, the viable count approach merely provides a broad overview of a population that is intrinsically different, and in all likelihood, it will be releasing diverse levels of cytokines and undergoing cell death at different rates at any given moment. In addition, uninfected cells within the well can mask the phenotype of infected cells with the phenotype of infected cells diluted by the presence of many uninfected cells in the same well with an opposing phenotype. In contrast, imaging flow cytometry here allowed the determination of bacterial number at the single cell level and permitted the correlation of bacterial load and Pyk2 status within each individual cell. It also allowed accurate determination of intracellular numbers of persistent, viable, non-culturable cells, which would be excluded using the viable count method. In addition, extracellular or attached bacteria could be discounted and uninfected macrophages within a well of infected cells could also be identified and analysed as a separate subpopulation. The high throughput nature of imaging flow cytometry provided sufficient data to enable trends to be determined within infected cells, such as Pyk2 status versus intracellular bacterial load. However, as with any approach there are limitations. Some bacteria may be lysed or non-viable, but their fluorescent signal may still be temporarily detectable. Additionally, the masks applied during analysis to determine cell size will never be 100% accurate for the entire population of cells, and there will always be cells that do not conform to the parameters set. However, applying this approach to phenotypically characterize a heterogenous population of infected cells based on their intracellular bacterial load is a significant step forward in studying infection at the level of the cell rather than the level of the well.

To date AIEC infection has been poorly understood relative to other pathogens, it’s paucity of virulence factors rendering it difficult to study via traditional microbiological approaches of mutation and testing. Its virulence has been attributed to its unique ability to survive and rapidly replicate within macrophages, while inducing a strong inflammatory response in the form of TNFα release. Here we focused on the AIEC-host interaction through high throughput imaging and phenotyping of infected cells populations and studying them in the context of Pyk2, a known susceptibility locus for development of IBD. Our approach enabled the removal of phenotypic noise introduced by the presence of many uninfected cells within infected cell populations, giving a clearer picture of the AIEC-macrophage relationship. Our results identified a crucial role for Pyk2 in facilitating AIEC uptake by macrophages, as previously reported for other pathogens, but remarkably also showed for the first time a critical role for Pyk2 in facilitating AIEC intra-macrophage replication and TNFα release. We also demonstrated how pharmaceutical intervention to block Pyk2 function could block AIEC intracellular replication and subsequent TNFα release, identifying a potential pathway towards an intervention strategy for CD patients where AIEC are dominant in the intestinal microbiome. The importance of these findings was underlined with reproduction of these phenotypes with clinical isolates of AIEC, while the relevance for other enteric pathogens was also shown through Pyk2 inhibition also significantly altering the course of *Salmonella* Typhimurium infection.

---

**Supplementary Fig. S1.**
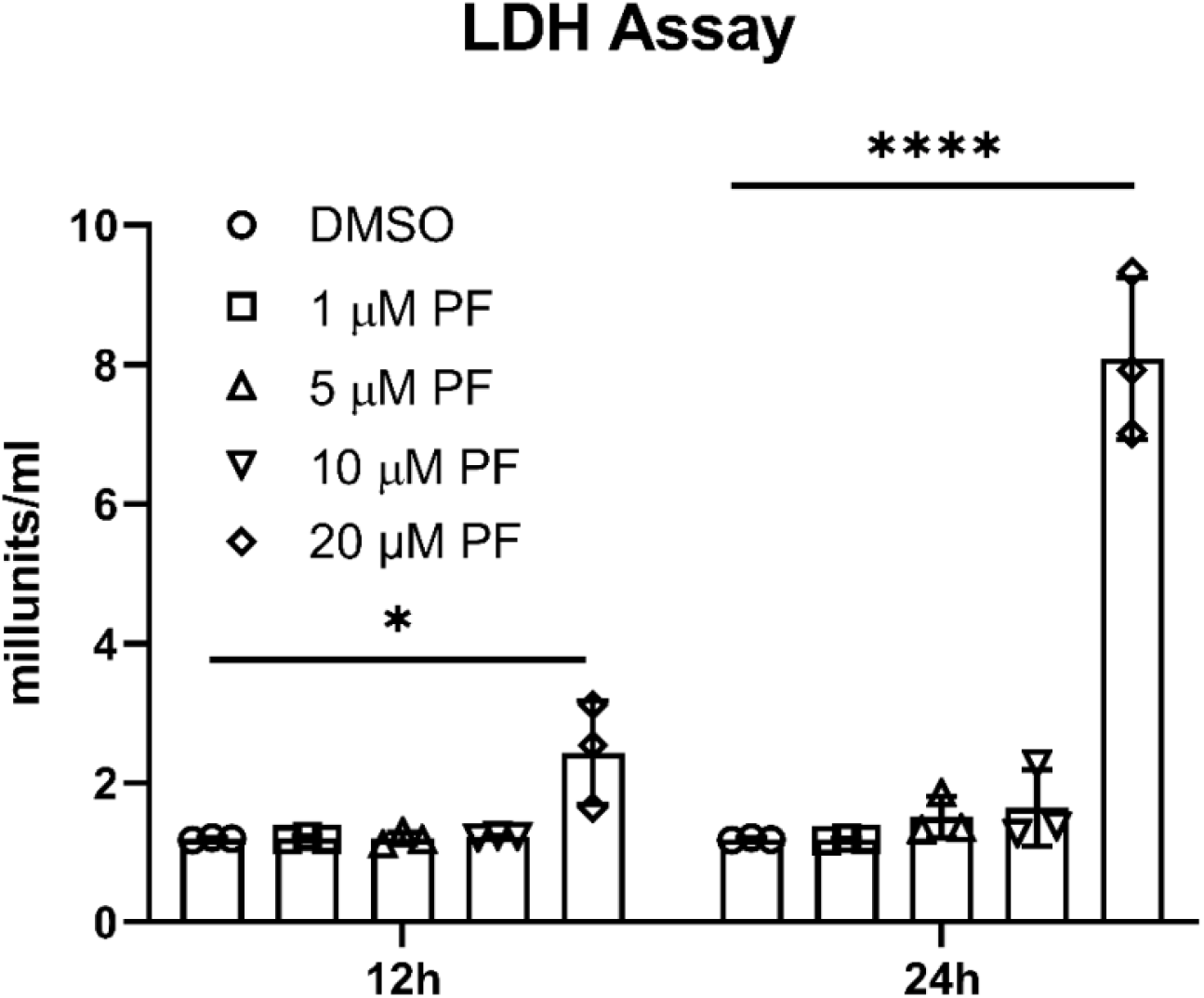
Low concentrations of Pyk2 inhibitor PF-431396 hydrate had no effect on cell toxicity. LDH activity assays were conducted on supernatants collected from RAW 264.7 cells following infection with LF82 and treatment with the Pyk2 inhibitor PF-431396 hydrate. LDH activity is reported as nmol/min/mL = milliunit/mL. Statistical analysis was conducted using a two-way ANOVA (ns, not significant, *p < 0.05, **p < 0.01, ***p < 0.001, ****p < 0.0001). Data are representative of three independent biological replicates.

**Supplementary Fig. S2:**
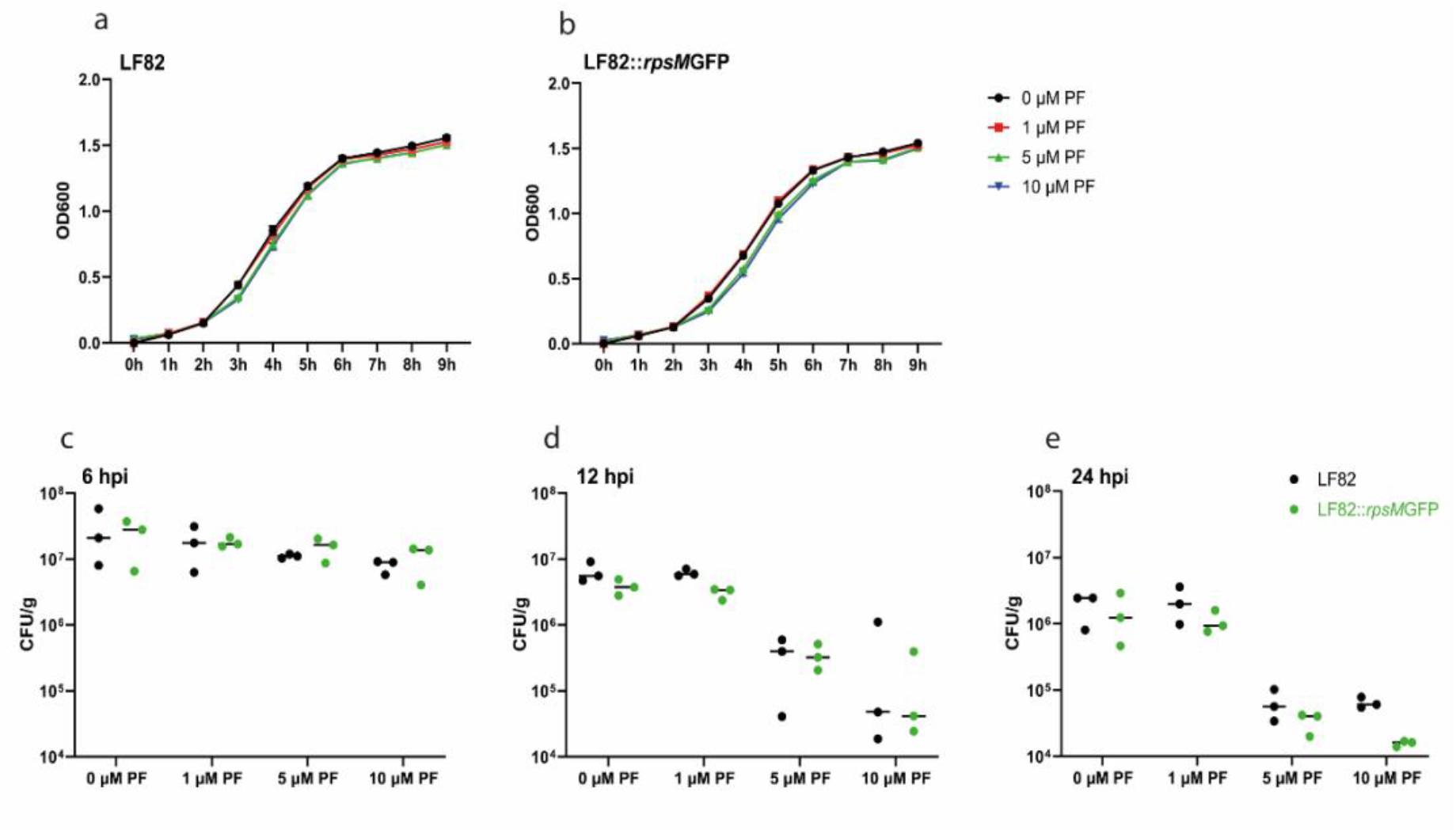
Transformation with *rpsM*GFP has no effect on growth or intracellular replication of LF82 in the presence or absence of PF-431396 hydrate. LF82 was transformed with *rpsM*GFP. Growth of LF82 (**a**) and LF82::*rpsM*GFP (**b**) was not affected by addition of increasing concentrations of PF-431396 hydrate. Intracellular replication of LF82 and LF82::*rpsM*GFP at 0 μM, 1 μM, 5 μM and 10 μM of PF-431396 hydrate was examined via viable colony counts at 6 (**c**), 12 (**d**) and 24 (**e**) hpi. Statistical analysis was conducted using a two-way ANOVA (ns, not significant, \**p* < 0.05, **p < 0.01, \*\*\**p* < 0.001, \*\*\*\**p* < 0.0001). Data (**c** – **e**) are representative of three independent biological replicates

**Supplementary Fig. S3:**
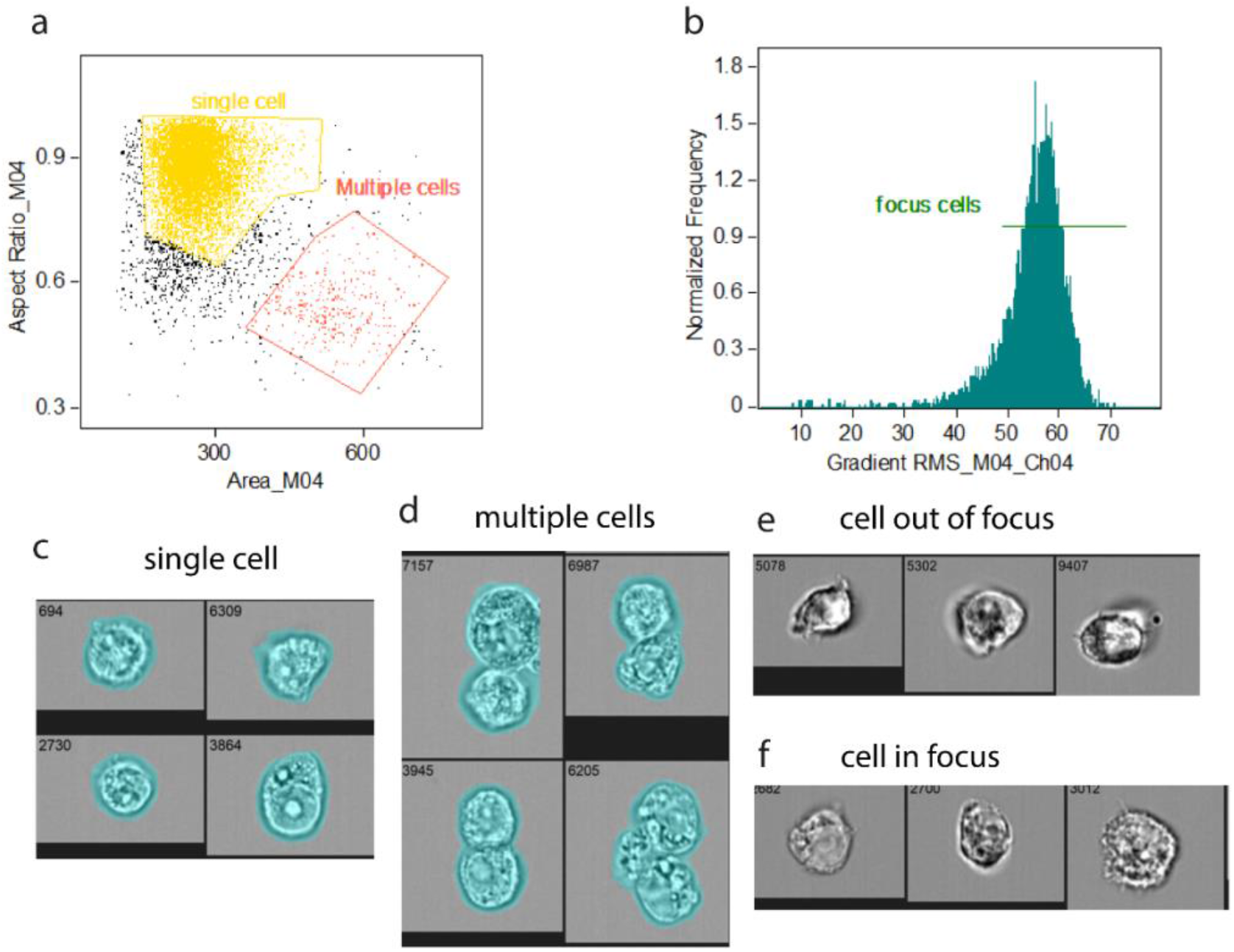
IDEAS analysis of LF82::*rpsM*GFP infected cells. RAW 264.7 were infected with LF82::*rpsM*GFP (MOI 100) for 1 h and analysed by imaging flow cytometry at 6 and 12 hpi. Single-cell population was defined by Area/Aspect ratio dot plot (**a**) and objects in best focus were gated as those events with gradient RMS values greater than 50 (**b**). Examples of cells that were included and excluded by the gating strategy; representative single cells (**c**), multiple cells (**d**), cells out of focus (**e)**; RMS value less than 50 as in [**b**]) and cells in focus (**f**; RMS value greater than 50 as in [**b**]).

**Supplementary Fig. S4:**
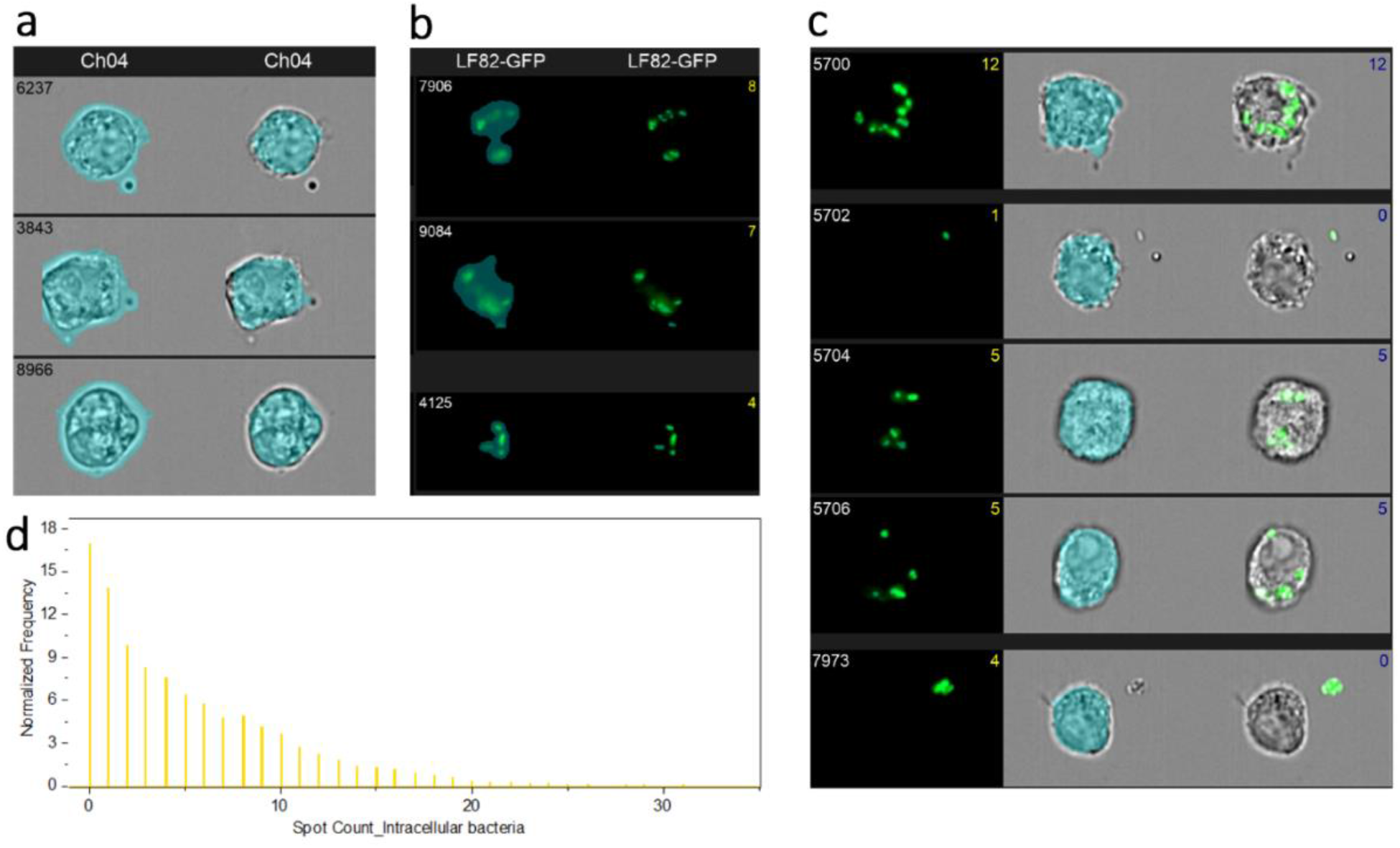
IDEAS intracellular and spot count masks. Intracellular bacterial localisation was measured by creating an intracellular mask (**a**). The spot count feature was used to quantify fluorescent spots identified using spot mask (computer code: Intensity (Peak[Spot{M02, LF82-GFP, Bright, 3.5, 3, 1}, LF82-GFP, Bright, 1]) (**b**). The spot mask in conjunction with the Erode mask was used to create an intracellular fluorescence count (**c**; computer code: Spot count-Intensity (Peak[Spot{M02, LF82-GFP, Bright, 3.5, 3, 1}, LF82-GFP, Bright, 1]) And AdaptiveErode (M04, Bright Field, 87, LF82-GFP, 80-4095). Quantitative distribution of GFP positive, single focused cells (RMS > 50) inside LF82-infected cells (**d).**

